# Modification of Death receptor-mediated cell death in osteosarcoma cell lines using epigenetic modifiers

**DOI:** 10.64898/2025.12.10.693227

**Authors:** Habitha Sri Prabhakaran, Neil Alan Cross

## Abstract

Osteosarcoma is a bone malignancy most prevalent in adolescent populations. Despite advances in multi-agent chemotherapies and surgical procedures, the survival rate remains <30% for patients with metastasis. Tumour necrosis factor-related apoptosis inducing ligand (TRAIL) is a promising anti-cancer agent, but when used as a single-agent, TRAIL-resistance occurs. Anti-Death receptor-4 and −5 agonistic mAbs are TRAIL mimetics proven to have higher specificity towards DR4 and DR5 receptor thus avoiding decoy receptors. However, their efficacy is also limited due to rapid development of resistance. In this study, the epigenetic modifiers Trichostatin A, GSK343 and BIX-01294 were explored in combination with an anti-DR5 agonistic mAb (DR5 mAb) to evaluate their anti-cancer efficacy in osteosarcoma cell lines Saos-2 and MG63. These cell lines are known to be insensitive TRAIL-induced apoptosis due to overexpression of membrane-bound and soluble decoy receptors to TRAIL, but are more sensitive to DR5 mAbs. Trichostatin A, a Histone deacetylase inhibitor (HDAC^i^) demonstrated a synergistic effect in sensitising osteosarcoma cells to DR5 mAb. Conversely, inhibitors of the Histone methyltransferase GSK343 (EZH2^i^) and BIX-01294 (G9a^i^) in combination with DR5 mAb had an antagonistic effect on apoptosis induction. The drug combinations were also evaluated in 3D alginate tumour spheroids, and both GSK343 and BIX-01294 inhibited DR5 mAb-mediated apoptosis.

## Introduction

Osteosarcoma is a primary bone malignancy with the most common prevalence in adolescent populations presented with extreme metastases to lung. They are characterized by the deposition of immature osteoid on bone by tumour cells (Cole et al., 2022). This malignancy has a unique epidemiological feature that vary by age, increased occurrence of axial tumours and metastatic disease and worse 5-year survival is found in older patients; and is the third most common type of cancer among children and adolescents after leukaemia and lymphoma. The incidence is noted to be approximately 6 cases per million in children under the age of 15. The most common locations of incidence being distal femur, proximal tibia, and proximal humerus in the mentioned order but in older patients the disease may develop in other bone regions (Cole et al., 2022). Current successful treatment options available for osteosarcoma are surgical resection followed be systemic chemotherapy regimens and adjuvant radiotherapy with a long-term impact on patient’s quality of life (Belayneh et al., 2021). The backbone of osteosarcoma treatment especially for younger patients is the chemotherapeutic MAP regimen: high-dose methotrexate (HD-MTX), doxorubicin (Adriamycin), cisplatin. This combination may be combined with a fourth regimen ifosfamide in higher-risk cases or non-responders for both neoadjuvant and adjuvant cases, achieving 5-year survival rate in approximately 70% of patients with localised disease but only 5-30% in cases with metastasis (Beird et al., 2022). Alternative regimens may be followed by substituting or adding drugs such as ifosfamide, carboplatin or etoposide in in cases where HD-MTX is not tolerated (Hu, et al., 2022). Despite the advances in multi-agent chemotherapies and surgical procedures, the survival rate remains unchanged over the years. Late-stage diagnosis and prevalence of high-grade osteosarcoma in adolescent patients’ needs to be addressed with specifically targeting therapeutic agents.

Tumour necrosis factor-related apoptosis inducing ligand (TRAIL) is a potential anti-cancer strategy as it selectively targets transformed cells and induces apoptosis without targeting non-transformed cells. This mechanism is carried out by TRAIL binding to the Death Receptor-4 and −5 (DR4 and DR5), inducing apoptosis through both intrinsic and extrinsic pathways, activating caspase cascades. Unfortunately, TRAIL also binds to decoy receptors such as DcR1, DcR2 and also the soluble decoy receptor osteoprotegerin (OPG), which halts the induction of apoptosis especially in osteosarcoma cell lines (Locklin, et al., 2007). Furthermore, bone marrow stromal cell-derived OPG may also protect tumour cells from TRAIL in a paracrine manner (Nyambo et al, 2004). Therapeutic monoclonal antibodies are made to overcome this resistance mechanism by binding specifically to DR4 and DR5, this can effectively bypass the decoy receptors and induce apoptosis in malignant cells effectively (Locklin et al., 2007). However, TRAIL and agonists of DR4 (von Pawel, et al., 2014) and DR5 (Forero, et al., 2017) in clinical trials have revealed limited anti-tumour efficacy due to the rapid resistance mechanism exhibited by cancer cells and poor pharmacokinetics observed (Bellail, et al., 2009). New generation death receptor agonists such as ozekibart (INBRX-109) also showed modest efficacy and did not have significant tumour shrinkage in clinical trials (Subbiah, et al., 2023). TRAIL is also known to activate non-apoptotic pathways in cancer cells including some survival pathways that may aid in immune surveillance, non-canonical signalling and tumour microenvironment modulation (Alves et al., 2020). The potential of TRAIL and rhTRAIL as a single agent is shown to be limited in anti-cancer treatment. Ongoing research focuses on finding TRAIL-sensitising agents by silencing or modifying the resistance mechanisms exhibited in response to TRAIL treatment.

One among such evaluated agents that has shown potential are epigenetic modifiers, which induce reversible changes in gene expression without altering the DNA sequence. Their mechanisms of action are include histone modification, modifying DNA methylation and chromatin structure therefore reprogramming cancer cells by either re-activating tumour suppressor gene (TSG) expression or by silencing oncogenes responsible for tumour growth promotion (Cheng, et al,. 2019, Lu, et al, 2020). Inhibitors of Histone Methyltransferase such as EZH2 (Enhancer of Zeste homologue 2) and G9a act in-part by inhibiting methylation of histones, therefore reactivating tumour suppressor genes, resulting in sensitisation of cancer cells when combined with other anti-cancer drugs. Histone modifiers such as HDAC^i^ (Histone Deacetylase inhibitors) modify histone acetylation which alters chromatin accessibility and gene transcription and can result in induction of apoptosis in cancer cells, inhibition of proliferation and regulation of cell-cycle (Cheng, et al., 2019, Lu, et al, 2020). The epigenetic modifiers used in the study to sensitise osteosarcoma cells to agonistic DR5 mAb were Trichostatin A, GSK343 and BIX01294.

Trichostatin A (TSA), a HDAC inhibitor that loosens the chromatin structure, leading to altered gene expression (putatively activating TSGs) which promotes cell cycle arrest and apoptosis in cancer cells. It arrests cell cycle at G0/G1 phase and upregulates pro-apoptotic proteins such as p53, downregulates p21 to reduce proliferation; also takes part in downregulating anti-apoptotic proteins such as Bcl-2, survivin and cFLIP, inducing the cells towards apoptosis (Cheng, et al., 2019). TSA and other HDAC^i^ has been proven to sensitise many cancer cells such as breast (Wong, et al., 2022), myeloma (Arhoma et al 2017a, Fandy, et al., 2006), ovarian (Park et al., 2009) and gastric (Li, et al., 2016) to TRAIL by down regulation of anti-apoptotic proteins, upregulation of death receptors, inhibition of survival pathways and activation of caspase cascades (Cheng, et al., 2019).

GSK343, a histone methyltransferase inhibitor, which reduces trimethylation of H3K27 by selectively inhibiting EZH2, resulting in reduced proliferation, reversal of EMT (Epithelial-to-Mesenchymal transition), upregulation of tumour suppressor genes and cell-cycle arrest at G0/G1 phase (Arhoma, et al., 2017b). GSK343 was shown to sensitise cancer cells such as multiple myeloma (Arhoma, et al., 2021), glioma (Yu, et al., 2017) and colorectal (Ying, et al., 2018) to TRAIL and rhTRAIL. In multiple myeloma, the cells are sensitised to TRAIL by GSK343 through upregulation of death receptors, activation of caspase cascades and reversal of TRAIL resistance mechanisms in myeloma (Arhoma, et al., 2021). In glioma cells, the mechanism of sensitisation happens through inhibition of EZH2 leading to reversal of EMT and suppressing cells with cancer stem cell phenotype (Yu, et al., 2017); also, by modulating canonical and non-canonical pathways such as NF-κβ and IB pathways (Scuderi, et al., 2022). In colorectal RKO cells, sensitivity to TRAIL was obtained through cell cycle arrest at G0/G1 phase (Ying, et al., 2018).

BIX-01294, a histone methyltransferase inhibitor, targets G9a/EHMT2 thereby reducing H3K9 methylation. This results in induction of apoptosis, proliferation inhibition and disruption of survival pathways in various cancers such as lung adenocarcinoma (Kim, et al., 2021), renal carcinoma (Woo, et al., 2018) and hepatocellular carcinoma (Namgung, et al., 2019) and multiple myeloma (Arhoma, A et al., 2017b) by downregulating anti-apoptotic pathways and metabolic pathways. BIX-01294 sensitises cancer cells to TRAIL and rhTRAIL by downregulating surviving and XIAP, upregulating DR5 alone and inducing apoptosis independent of G9a activity (Woo, et al., 2018, Namgung, et al., 2019).

The research focuses on evaluating the efficacy of above-mentioned epigenetic modifiers in sensitising osteosarcoma cells (Saos-2 and MG-63) to DR5 mAb. These cell lines are known for their resistance towards Cisplatin and Doxorubicin, and one of their main mechanisms of resistance seems to be activation of survival pathways such as PI3K/AKT/mTOR (Low, et al., 2022, Gallego, et al., 2022). Both osteosarcoma cells are known to highly express OPG, a soluble decoy receptor that prevents binding of TRAIL making them insensitive to TRAIL-based therapies (Locklin et al., 2007). Hence, DR5 mAb was tested in combination with the epigenetic modifiers Trichostatin A (TSA), GSK343 and BIX-01294 to evaluate their potential synergistic effects in inducing apoptosis in osteosarcoma cell lines.

In this study, the general hypothesis tested was that epigenetic modifiers sensitise osteosarcoma cells to DR5 mAb. To test this hypothesis, the aims were to a) determine if osteosarcoma cells Saos-2 and MG63 are sensitive to Anti-DR5 agonistic mAb and epigenetic modifiers alone, b) assess whether TSA, GSK343 and BIX-01294 can sensitise osteosarcoma cells to DR5 mAb and to establish whether any synergistic effect is observed in Saos-2 and MG63 cells and c) assess whether combination treatment has the same effect in the 3D cell culture environment.

## Materials and Methods

### Cell culture and maintenance

Saos-2 and MG63, two osteosarcoma cell lines were obtained from American Type Culture Collection (ATCC) (Manassas, VA, USA). Saos-2, an osteosarcoma cell line with epithelial morphology was obtained from the bone of a 11-year-old, white female osteosarcoma patient. MG63, a cell line with fibroblast morphology was obtained from a 14-year-old, white male patient with osteosarcoma. The cell lines were maintained in MEM Alpha media and were supplemented with 10% FCS (Foetal calf serum), 2mM L-glutamine, 1% non-essential amino acids, 100 U/mL penicillin and 100mg/mL streptomycin. Cell cultures were maintained in incubator at 37°C and 5% CO_2_ (Phillips, et al., 2019). For 2D cell culture, 96 well plates were seeded with 2.5 × 10^3^ cells/ well for both MG63 and Saos-2 cell lines and were allowed to adhere to the well plate overnight in the incubator at 37°C in 5% CO_2_ prior to treatments for 24-hour treatment analysis.

### 3D Alginate bead culture

Cells density of 1× 10^6^ cells/ ml of osteosarcoma cells were resuspended in 1.2% sodium alginate prepared in saline 0.15M NaCl_2_. The cells resuspended in alginate solution were dropped into 0.2M CaCl_2_ using a 100µl pipette and allowed to incubate for few minutes at 37°C. The formed beads were washed thrice with 0.15M NaCl_2_. The beads were then suspended in MEM Alpha media supplemented with 10% FCS (Foetal calf serum), 2mM L-glutamine, 1% non-essential amino acids, 100 U/mL penicillin and 100mg/mL streptomycin in a T-25 flask and kept upright to avoid beads adhering to the surface. Each 1ml of cell suspension generated approximately 30 beads, each bead containing approximately 3.3 × 10^4^ cells. One bead per well was added and was treated the same day to avoid adherence or breaking of beads. (Phillips et al, 2019, Arhoma A et al, 2017a).

### Treatment with DR5 mAb and epigenetic modifiers in both 2D cell culture and 3D cell culture

To treat osteosarcoma cells, Anti-DR5 (MAB631) (R&D, Abingdon, Oxfordshire, UK was initially prepared as a stock of 2µg/ml in PBS and were stored as 10 µl aliquots. These aliquots were then used to prepare 5-fold dilutions with working concentrations ranging from 0-500 ng/ml (0, 1.6, 8, 40, 200, 500 ng/ml) in MEM Alpha media. For vehicle control, the 0.05% PBS in 1ml media was prepared from 1x PBS. TSA (Sigma, Poole, Dorset), UK) was initially prepared as stock of 10mM in ethanol and was stored in 10 µl aliquots. These aliquots were then used to prepare 5-fold dilutions with working concentrations ranging from 0-10 µM (0, 0.08, 0.4, 2, 10 µM) in MEM Alpha media using serial dilution technique. For vehicle control, the 0.05% ethanol was prepared by dissolving 10µl of neat ethanol in 1ml media. BIX-01294 (Sigma) and GSK343 (Sigma) were prepared into stock solutions of 10mM in DMSO and was stored in 10 µl aliquots. The aliquots were used to prepare working concentrations in 5-fold dilutions ranging from 0-10 µM (0, 0.08, 0.4, 2, 10 µM) in media. For vehicle control, the 0.05% of DMSO was prepared by dissolving 10µl DMSO in 1ml media.

Cells were treated with DR5 mAb alone and epigenetic modifiers (Trichostatin A, GSK343 and BIX01294) alone in their five-fold dilutions to determine their percentage of apoptosis as a single agent. Doses of 0-500 ng/ml of Anti-DR5 and doses of 0-10 µM of Trichostatin A, GSK343 or BIX-01294 were used to treat both osteosarcoma cell lines. To determine the effect of combination treatment, lowest dose that induced significant apoptosis were chosen; and for drugs with no significant apoptosis maximum dose were chosen. The drug doses chosen were DR5 mAb (200 and 500 ng/ml), Trichostatin A (2µM), GSK343 (10µM) and BIX-01294 (10µM). Control cells were treated with 0.05% PBS for DR5 mAb; 0.05% of DMSO for GSK343 and BIX-01294; 0.05% of ethanol for Trichostatin A.

### Assessment of apoptosis through Hoechst 33342 and PI staining

Assessment of apoptosis was done by staining both 2D and 3D cells with 1 µl of 10mg/mL Hoechst 33342 (Sigma) and 10 µl of 10mg/mL Propidium Iodide (Sigma) at 37°C in the incubator for 30 mins. Cells were visualised and captured using EVOS microscope or an Olympus IX70 inverted microscope and analysed using Cell^F software. Apoptosis was determined manually by assessing the nuclear morphology. Triplicate images from each well with at least 200 cells were used to determine percentage of apoptosis. For combination treatments triplicate values from three independent experiments were considered to determine percentage of apoptosis.

### Statistical Analysis

For statistical analysis all the experiments were done in triplicates (n=3) and was calculated for mean and SD. Single treatments were analysed with Kruskal-Wallis test with Dunnett *post-hoc* test to determine the sensitivity at significant concentration and for combination treatments two-way ANOVA with Holm Sidak *post-hoc* test was done to find out synergistic significance between compared treatments using GraphPad Prism software version 10 (GraphPad Software Inc., CA, USA). SynergyFinder+ (Tang et al., 2022) was used to calculate the percentage of synergistic or antagonistic effect between compared treatments (single vs combination).

## Results

The sensitivity to Anti-DR5 agonistic mAb and the epigenetic modifiers Trichostatin A, BIX-01294 and GSK343 were assessed in Saos-2 and MG63 osteosarcoma cell lines using Hoechst 33342 and Propidium iodide staining (Figure. 1). Saos-2 and MG63 showed significant induction of apoptosis using DR5 mAb at 200ng/ml and 500ng/ml (Figure 1). Of the epigenetic modifiers, Trichostatin A potently induced apoptosis in both Saos-2 and MG63 cell lines at 10µM. Induction of apoptosis was noticeably higher in Saos-2 (40%) when treated with GSK343 than in MG63 (4%). For BIX-01294 treatment, Saos-2 cells were sensitive although to a lesser extent than MG63 cells. All epigenetic modifiers were observed to show sensitivity in both osteosarcoma cell lines at their highest treated concentration of 10µM except for GSK343 in MG63.

**Figure. 1.**
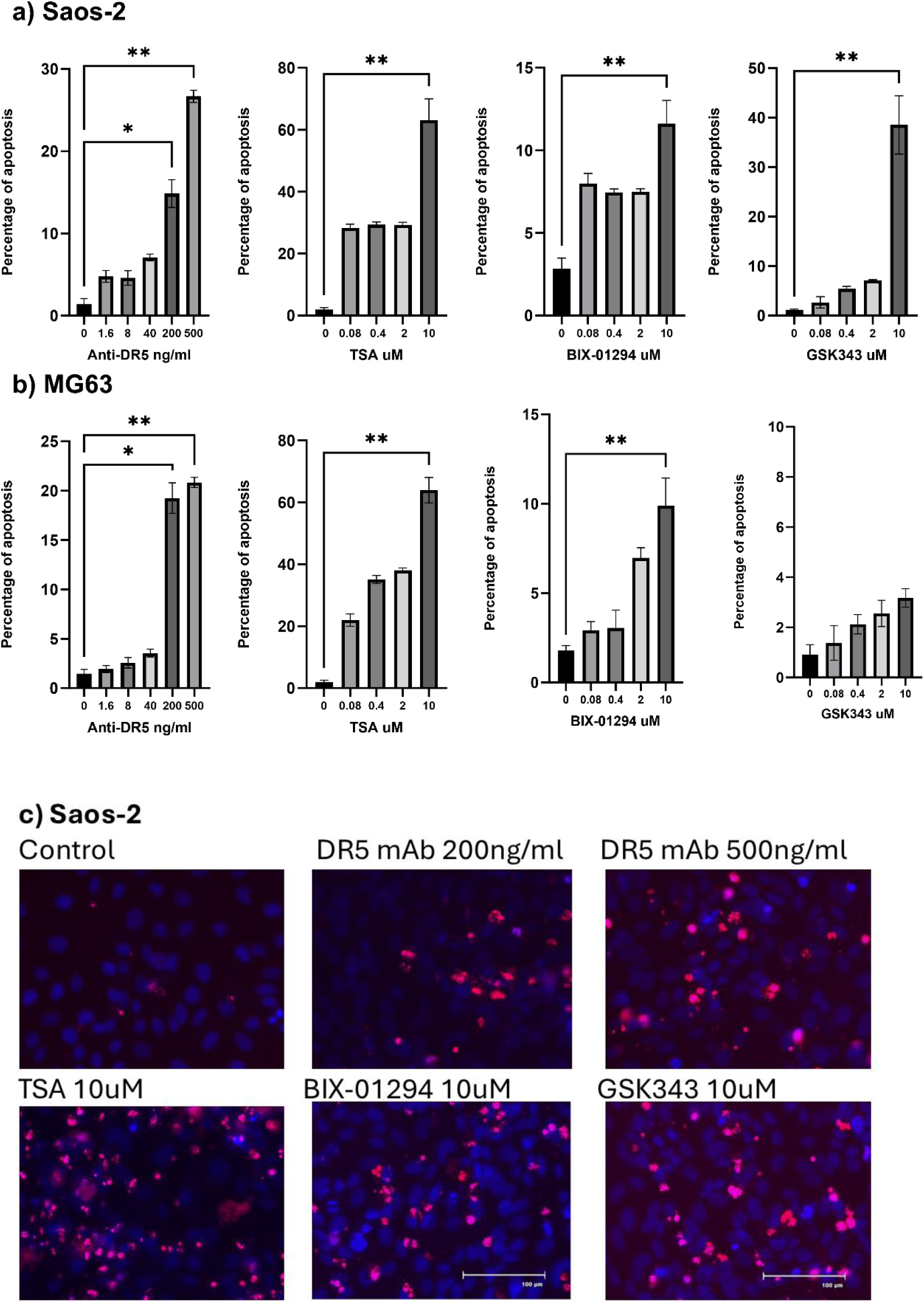
Analysis of apoptosis in response to single agent treatment was done using Hoechst 33342 and PI staining in both A) Saos-2 and B) MG63 osteosarcoma cell lines following 24 hours stimulation with Anti-DR5 (0-500 ng/ml), Trichostatin A (0-10 µM), BIX01294 (0-10 µM), GSK343 (0-10 µM). A) Statistical analysis was done using Kruskal-Wallis test with Dunnett post hoc test with n=3 and values are calculated as Mean +/− SD. * indicates p value < 0.05; ** indicates p value < 0.005. C) Representative Hoechst 33342 and PI images done on Saos-2 cells in response to vehicle control, Anti-DR5 (200 & 500 ng/ml), Trichostatin A (2 µM), BIX01294 (10 µM), GSK343 (10 µM). Condensed and/or fragmented nuclei stained blue are early apoptotic cells and cells that are stained red (PI positive) are late apoptotic cells and a small number of necrotic cells (red, not condensed/fragmented).

### Assessment of apoptosis in response to combination treatment in 2D cells through Hoechst 33342 and PI staining

Combination treatment was conducted to determine whether epigenetic modifiers can sensitise osteosarcoma cells to DR5-mAb. Two concentrations of DR5 mAb were selected: 200 ng/ml, which was proven to induce apoptosis of <20% in both Saos-2 and MG63 cells and 500 ng/ml (<30% apoptosis) to examine the synergistic effect when combined with epigenetic modifiers. The osteosarcoma cells were stimulated with a concentration of epigenetic modifiers that induced <40% apoptosis: Trichostatin A (2µM), BIX01294 (10µM) and GSK343 (10µM), were selected for combination treatment analysis. Cells were assessed using Hoechst 33342 and PI staining. Drug interaction analysis was performed using SynergyFinder+ to confirm synergistic effect in response to dual treatment through Bliss score.

The osteosarcoma cells lines analysed show a high sensitivity to Trichostatin A (TSA) at 2µM in Saos-2 cells. Combination treatment of TSA with Anti-DR5 at 200 and 500 ng/ml were showed synergy in both cell lines (Figure. 2 and 3); in comparison to both the doses of DR5 mAb and TSA individually. The Combination Index (CI) was consistently < 1 for the dual treatment of DR5 mAb and TSA in both cell lines. The synergistic effect was noted to increase as the DR5 mAb dose increases (Figure 3).

**Figure. 2.**
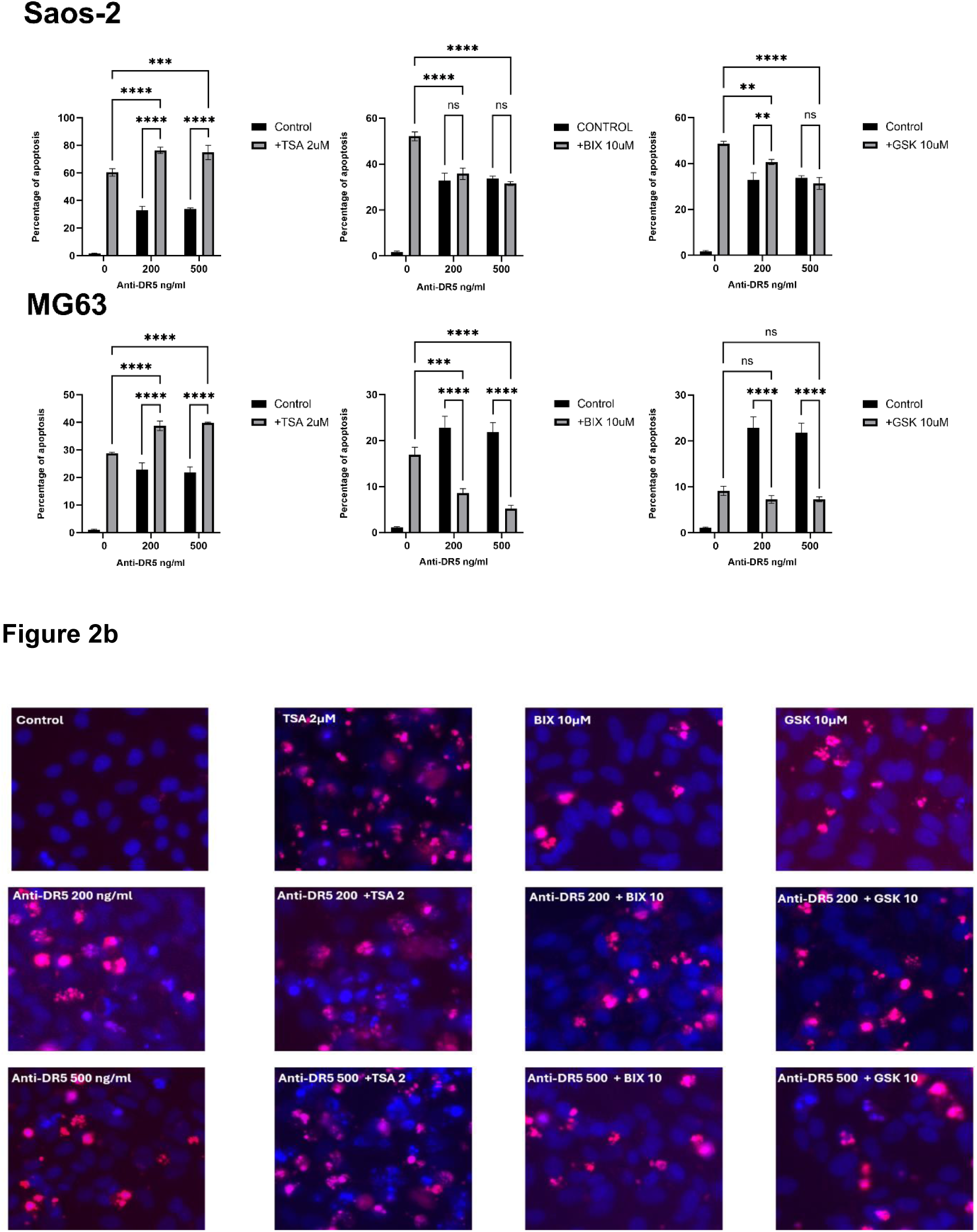
Analysis of apoptosis in response to combination treatment was performed using Hoechst 33342 and PI staining in Saos-2 and MG63 osteosarcoma 2D cell cultures following 24-hour stimulation with epigenetic modifiers Trichostatin A (2 µM), BIX-01294 (10 µM) or GSK343 (10 µM) in the presence or absence of DR5 mAb (200 & 500 ng/ml). A) Data were presented in Mean +/− SD with n=3; statistics performed using two-way ANOVA with Holm-Sidak post hoc test. *** indicates p value > 0.0001; **** indicates p value < 0.0001. Synergistic enhancement is observed with TSA (see figure 3), whilst antagonism of DR5 mAb-induced cell death is observed with BIX-10294 and GSK343 (see figure 3). B) Representative images of Hoechst 33342 and PI staining performed on the Saos-2 osteosarcoma cell line in response to vehicle control, Anti-DR5 (200 & 500 ng/ml), Trichostatin A (2 µM), BIX-01294 (10 µM), GSK343 (10 µM), Anti-DR5 + TSA, Anti-DR5 + BIX, Anti-DR5 + GSK. Condensed nuclei stained blue (H positive) are early apoptotic cells and cells that are stained red (PI positive) are late apoptotic cells.

**Figure 3.**
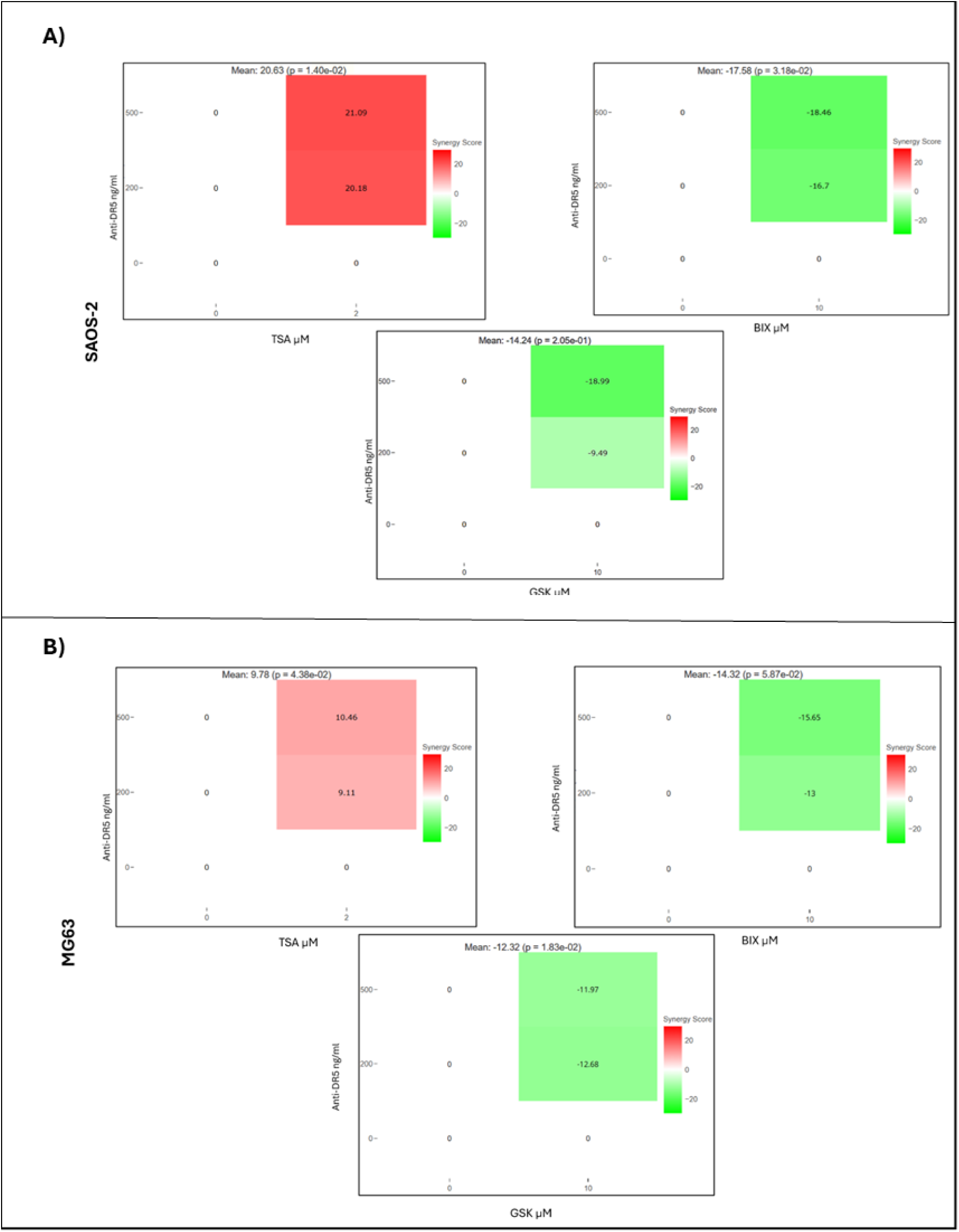
Representative outputs from SynergyFinder+ software used to analyse Bliss score to calculate synergistic or antagonistic effect obtained when DR5 mAb is combined with epigenetic inhibitors (TSA, GSK, BIX) in Saos-2 (A) and MG63 (B) 2D cells. Red indicates the synergistic effect in response to combination treatment. Green indicates the antagonistic effect in response to combination treatment. The mean value here represents the overall percentage of synergistic (+), or antagonistic (-) effect observed in response to combination treatment.

In Saos-2 cells, BIX-01294 had a notable antagonistic effect on induction of apoptosis when combined with DR5 mAb, with a decrease in apoptotic induction as the DR5 mAb dose increases (Figure 2 and 3) and the antagonistic effect was also noted in MG63 cells (Figure. 2 and 3). GSK343 also had an antagonistic effect in apoptosis induction in Saos-2 and MG63 cells when combined with varying doses of DR5 mAb (Figure. 2 and 3). Both the drug combinations had a dose-dependent antagonistic effect in response to DR5 mAb.

### Assessment of apoptosis in response to combination treatment in 3D Alginate spheroids by Hoechst 33342 and PI staining

To evaluate the combined effect of epigenetic modifiers and DR5 mAb, the osteosarcoma cells were encapsulated in 3D alginate beads and allowed to form spheroids and were exposed to combined treatment with TSA, GSK343 and BI-01294 with DR5 mAb. The responses were observed through Hoechst 33342 and PI staining. Drug interaction analysis was performed using SynergyFinder+ to confirm synergistic effect in response to dual treatment through Bliss score.

The drug interaction analysis of osteosarcoma cells in 3D spheroids revealed that GSK343 had an antagonistic effect when combined with DR5 mAb with a decrease in apoptotic induction as the DR5 mAb dose increases in both cell lines (Figure 4). In Saos-2 cells and MG63 cells, the antagonistic effect was noted for the treatment with GSK343 (Figure 4). TSA potentially killed Saos-2 cells more potently in 3D cell culture than in 2D cell culture, and DR5 mAb partially reversed this. In MG-63, there was no synergistic induction or antagonism noted in 3D cell culture (Figure 4).

**Figure. 4.**
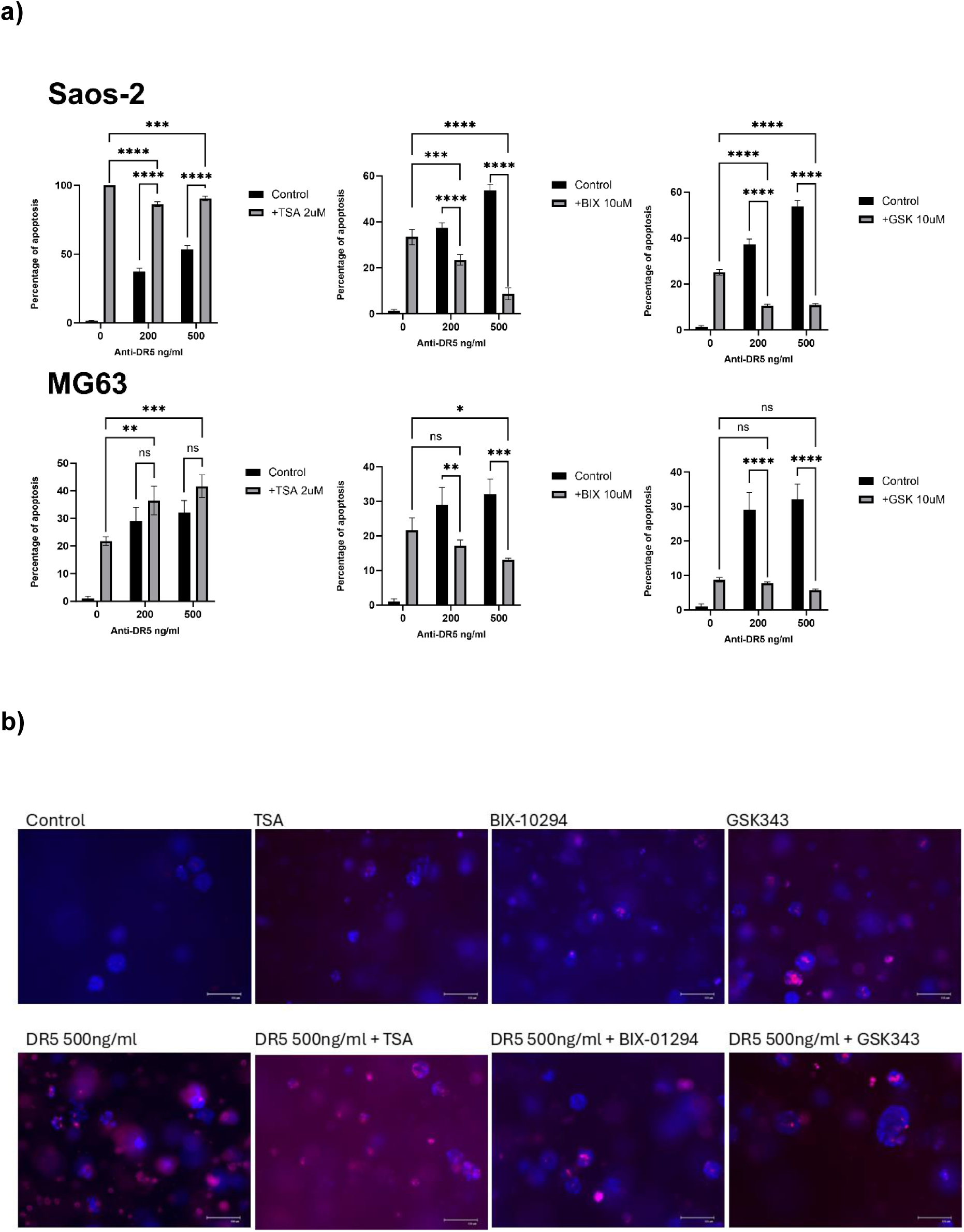
Analysis of apoptosis in response to combination treatment was done using Hoechst 33342 and PI staining in Saos-2 and MG63 osteosarcoma 3D alginate beads following 24-hour stimulation with Epigenetic modifiers Trichostatin A (2 µM), BIX01294 (10 µM) or GSK343 (10 µM) in the presence or absence of Anti-DR5 (200 & 500 ng/ml); to count percentage of apoptosis 10-15 colonies were counted from each treated beads. A) Data were presented in Mean +/− SD with n=3; statistics performed using two-way ANOVA with Holm-Sidak post hoc test. * indicates p value < 0.05; ** indicates p value < 0.005; *** indicates p value > 0.0001; **** indicates p value < 0.0001. TSA potently induced cell death in Saos-2, and DR5 mAb partially reversed this. DR5 mAb partially reversed GSK343 and BIX-12494-mediated effects also in Saos-2 B) Representative images of Hoechst 33342 and PI staining performed on Saos-2 osteosarcoma cell line in response to vehicle control, DR5 mAb (500 ng/ml), Trichostatin A (2 µM), BIX01294 (10 µM), GSK343 (10 µM), or combination with DR5 mAb (500ng/ml). Condensed nuclei stained blue (H positive) are early apoptotic cells and cells that are stained red (PI positive) are late apoptotic cells or dead cells.

## Discussion

This study aimed to evaluate the effectiveness of TRAIL-based drug, DR5 mAb against osteosarcoma cell lines Saos-2 and MG63 in inducing apoptosis as a single-agent. They were also analysed in combination with the epigenetic modifiers TSA, GSK343 and BIX01294. The study shows that DR5 mAb in combination with TSA results in a synergistic effect in Saos-2; and in MG63 to a lesser extent. GSK343 and BIX01294 were observed to have antagonistic effect when combined with DR5 mAb in both Saos-2 and MG63 cells as the DR5 mAb dose increases. The synergistic and antagonistic effects observed in response to combination treatments analysed showed similar effects in 3D culture conditions of both Saos-2 and MG63, although in Saos-2, TSA proved lethal, and responses were partially revered by DR5 mAb.

### TSA enhances TRAIL-induced apoptosis in osteosarcoma cell lines

In this study, TSA, a HDAC inhibitor was evaluated on Saos-2 and MG63 osteosarcoma cell lines to assess their effectiveness as single-agents and in combination with DR5 mAb. The findings showed that TSA as a single-agent was proven to have significant induction of apoptosis in Saos-2 and MG63 (Figure 1). It also showed significant synergistic effect when combined with DR5 mAb in Saos-2; in MG63 the combination had a minimal effect in comparison to Saos-2 (Figure. 2).

Consistent with the findings HDAC^i^ have shown to exhibit promising effect in sensitising TRAIL in various cancers. In colorectal cancer, HDAC inhibitors shown to reverse TRAIL-resistance and induce apoptosis via intrinsic pathway and by upregulating DR5 (Cui, et al., 2022). The combination was noted to promote apoptosis in leukaemia and lymphoid malignancies by triggering apoptosis through death receptor and mitochondrial pathways (Rosato, et al., 2003). Combining class 1 HDAC inhibitors with TRAIL variants induced apoptosis in TRAIL-resistant colon cancer cells by upregulating DR 4/5 (Zhang, et al., 2019).

TSA synergises with TRAIL and TRAIL variants by downregulating c-FLIP and anti-apoptotic Bcl-2 proteins thereby removing the blocks on extrinsic and intrinsic pathways, inhibits survival pathways such as EGFR/ERK/FOXM1 reversing TRAIL resistance, activates caspase cascades (Sayers and Cross, 2014, Kong, et al., 2019, Wong, et al., 2022). TSA when combined with TRAIL or TRAIL variants have shown to sensitise various cancer cells. In gastric cancer, TSA showed promising synergic effect with TRAIL by enhancing the cells sensitivity through inhibition of ERK/FOXM1 pathway (Li, et al., 2016). In breast cancer, TSA in combination with other epigenetic drug such as zebularine and TRAIL significantly induced apoptosis in triple negative and other breast cancer subtypes, with minimal effect on normal cells showing target specificity towards cancer cells (Kong, et al., 2019, Wong, et al., 2022). Consistent with these studies, the osteosarcoma cells treated with TSA 2µM alone induced significant apoptosis-mediated cell death (Figure 1) and when combined with anti-DR5 (200 and 500 ng/ml) had a synergistic effect in inducing apoptosis compared to cells treated with DR5 mAb alone (Figure 2). In 3D spheroids, a more potent TSA response was observed which was partially reversed by DR5 mAb (Figure 4).

### GSK343 exhibits antagonistic interaction with DR5 mAb in osteosarcoma cells

EZH2 inhibitors targets EZH2, a catalytic subunit of Polycomb Repressive Complex 2 (PRC2) responsible for trimethylation of H3K27 leading to transcriptional repression. They have demonstrated anti-cancer efficacy in various cancer cells including osteosarcoma, as mono-therapeutics and in dual treatment with other epigenetic modifiers or chemotherapeutics. In colon cancer, EZH2 inhibitors in combination with DNMT inhibitors have shown to synergically upregulate tumour suppressor genes and inhibit proliferation (Tiedemann, et al., 2025). In small cell lung cancer, the EZH2 inhibitor was proven to restore chemosensitivity in pre-clinical models and dual inhibitor of EZH1/2 were observed to outperform in standard of care therapies (Han, et al., 2025). EZH2^i^ in combination with carboplatin was proven to be effective in enhancing the activity of carboplatin in aggressive variant of prostate cancer cells by reducing the expression of DNA repair genes and by upregulating expression of p53 dependent apoptosis aiding factors (Latarani, et al., 2025). In oesophageal adenocarcinoma, the EZH2 inhibitors were noted to be effective both as a single-agent and in combination with 5-FU; in combination the efficacy was observed as they reduced the IC50 of 5-FU in *in vitro* studies (Pickering, et al., 2022). EZH2i were observed to be effective as a mono-therapy agent in colorectal cancer, by directly suppressing cell proliferation and by shifting macrophage polarisation towards a tumour suppressive phenotype in tumour microenvironment (Li, et al., 2022).

EZH2 are found to be overexpressed in osteosarcoma cells associated with poor prognosis and aggressive tumour behaviour (De Carvalho, et al., 2012). In osteosarcoma cells, GSK343, a EZH2^i^ was proven to be effective by induction of apoptosis and autophagy, suppressing migration and invasion through down-regulation of oncogenic pathways such as c-Myc and FBP1 axis (Xiong, et al., 2016, Xiong, X et al., 2016). EZH2 inhibitors were proven to reduce the growth of tumour and decrease the metastases to lung, a major cause of death in osteosarcoma metastatic patients (Devarajan, et al., 2024). Consistent with this research outcome, the osteosarcoma cells treated with GSK343, was proven to induce significant apoptosis as a single agent at a dose of 10µM (Figure. 1).

EZH2i in combination with TRAIL-based therapies have proven to be effective in various cancer types. In myeloma cells, the combination of GSK343 with TRAIL was noted to increase TRAIL sensitivity in both TRAIL sensitive and TRAIL resistant cells by activating caspase-8 and caspase-9 pathways. But the sensitisation effect was not observed in quiescent cells or non-dividing cells limiting their effectiveness to proliferating cells (Arhoma, et al., 2021). EZH2^i^ (EPZ-6438 and PF-06821497) in combination with ONC201/TIC10, an impridone class anti-cancer drug upregulating DR5 was found to be effective across various cancer types such as glioma, breast cancer including triple-negative subtype etc. through enhancement of sensitivity towards ONC201, evidenced by increased apoptotic markers such as cleaved-PARP and by upregulating DR5 (Zhang, et al., 2021). There are no direct studies done to evaluate the efficacy of EZH2^i^ with TRAIL-based therapies in osteosarcoma. Here, the study was done to evaluate the combination’s anti-cancer efficacy in osteosarcoma cells and revealed a contradictory finding to other studies done on various cancer cells, GSK343 in combination with DR5 mAb had an antagonistic effect in comparison to GSK343 alone (Figure. 3). The antagonistic effect was noted to be increasing as the doses of Anti-DR5 increases (Figure. 3). The same antagonistic effect was observed in 3D spheroids in response to the treatment and the 3D spheroids help visualise the corelation between increase in DR5 mAb dose and increase in antagonistic effect (Figure. 4).

### BIX-01294 exhibits antagonistic interaction with DR5 mAb in osteosarcoma cells

G9a inhibitors target histone methyltransferase G9a, which catalyses mono- and di-methylation of H3K9, associated with repressed transcriptional activity. By inhibiting G9a, they promote tumour suppressor gene activation leading to tumour growth suppression. G9a inhibitors were established to have anti-cancer effect in various cancer types. In leukaemia cells *in vitro* studies, DCG066, a G9a inhibitor shown to dramatically reduce proliferation and self-renewal of acute myeloid leukaemia cells by gradually reducing Hox-A9 dependent transcription (Kondengaden, et al., 2016). In nasopharyngeal carcinoma, BIX-01294, a G9a inhibitor proved to suppress cell proliferation by inhibiting autophagic flux through inhibition of lysosomal cathepsin D activation (Li, et al., 2021). In castration resistant prostate cancer, CM-272, a dual inhibitor of G9a and DNMT1, caused G9a inhibition leading to decreased cell viability (Moreira-Silva, et al., 2022). SDS-347, a competitive G9a inhibitor, showed promising anti-cancer effect against various cancer cells, evidenced by reduction in H3K9me2 levels in a cell-based assay (Jan, et al., 2023). There are no studies do date evaluating the anti-cancer effect of G9a inhibitors in osteosarcoma cell lines. BIX-01294, a G9a inhibitor was used in this study to analyse their anti-cancer efficacy in osteosarcoma cells. The findings showed significant induction of apoptosis at a dose of 10µM in both Saos-2 and MG63 cells.

In many dual treatments, G9a inhibitors showed promising results in cancer treatments across various studies. UNC0638, a G9a inhibitor showed to sensitise osteosarcoma (U2OS) cells to etoposide (a DNA double strand break-inducing agent) and identified that this combination induced cell death independent of p53 (Agarwal, et al., 2016). In glioblastoma, BIX-01294 in combination with temozolomide sensitised tumour and stem-cell-like cells by enhancing apoptosis and autophagy evidenced through cleaved caspase-3, caspase-7 and PARP. These markers were also observed in resistant cells regardless of resistance and pluripotency markers such as NANOG, SOX2, CD133 (Ciechomska, et al., 2018).

BIX-01294 showed to enhance TRAIL-induced apoptosis in combination with TRAIL in renal carcinoma, breast carcinoma and lung carcinoma cells through downregulation of surviving, an anti-apoptotic protein found to be overexpressed in all the mentioned cancer cells and by upregulating DR5 (Woo, et al., 2018). In this study, a similar approach was taken to evaluate the TRAIL-sensitising effect of BIX-01294 on Saos-2 and MG63, as there are no available studies done on osteosarcoma cells. Contradictory to other studies done on various cancer types, BIX-01294 had an antagonistic effect when combined DR5 mAb in sensitising TRAIL to induce apoptosis. This was in direct correlation with increase in DR5 mAb doses. The same outcome was observed in 3D spheroids of the osteosarcoma cells, which help visualise the corelation between increase in Anti-DR5 dose and increase in antagonistic effect (Figure. 4).

### DR5 signalling associated with pro-survival and proliferative pathways

In line with the findings of antagonistic effect observed from the combinations GSK343 combined with DR5 mAb and BIX-01294 combined with DR5 mAb, recent studies have revealed that DR5 signalling has been associated with triggering of pro-survival and proliferative pathways such as NF-κβ, PI3K/Akt, MAPK and ERK1/2 which leads to a fraction of cell escaping the major apoptotic pathway of the DR5 signalling and proceeding down the survival and proliferation pathways leading to development of TRAIL resistance (Shlyakhtina, et al., 2017). Intriguingly, in paediatric bone cancers, AMG655, a DR5 agonist mAb induces apoptosis in TRAIL-sensitive cell lines but accelerates tumour growth in TRAIL-resistant cells via recruitment of RIPK1 (Brion, et al., 2022).

In conclusion, this study shows for the first time that TSA in combination with DR5 mAb had a promising synergistic effect in inducing apoptosis on tested osteosarcoma cells in both 2D cells but in 3D spheroids, responses were more potent in response to TSA alone, but DR5 mAb partially reversed cell death. The HMT^i^s GSK343 and BIX01294 had an antagonistic effect in on DR5 mAb apoptosis which corelates to DR5 mAb dose in both 2D cells and 3D spheroids of Saos-2 and MG63. The antagonistic effect can be correlated to the recent findings that DR5 signalling is also associated with activation of pro-survival and proliferation pathways. Further research is needed to investigate the mechanism behind the antagonistic effect of HMT^i^ in combination with DR5 mAb and to find out if the DR5 signalling is involved in activation of survival pathways in osteosarcoma cells.

## Acknowledgements

This work was completed as part of Habitha Sri Prabhakaran’s MSc research project. We thank the technicians for continued cell culture support throughout the project.

